# Oligonucleotide synthesis errors are a source of untoward variation in HDR-mediated gene editing

**DOI:** 10.1101/2025.07.08.663753

**Authors:** Stacia K Wyman, Zulema Romero, Seok-Jin Heo, Marian Navarrete, Netravathi Krishnappa, Donald B Kohn, David IK Martin, Mark C Walters, Dario Boffelli

## Abstract

Single-stranded oligonucleotides (ssODNs) are used as donor templates for therapeutic gene editing by CRISPR-Cas9 cleavage and homology-directed repair (HDR). Although ssODN sequence fidelity is critical to the safety and efficacy of editing, standard quality control methods cannot resolve individual nucleotide errors. By deep sequencing ssODNs from three manufacturers, and amplicons from edited hematopoietic stem/progenitor cells, we find that synthesis errors are present in all ssODNs tested at rates that vary more than two-fold among manufacturers, at positions that are dependent on sequence context. These synthesis errors are propagated into the genome by HDR at frequencies proportional to their abundance in the ssODN. In our sickle cell mutation correction protocol, the most prevalent SNEs are predicted to produce benign β-globin variants, while the less frequent frameshift deletions will generate β-thalassemia alleles. Current quality control standards are insufficient to detect these errors, and deep sequencing of ssODNs should be incorporated into regulatory submissions for clinical gene editing programs.

## Main

Single-nucleotide mutations that cause genetic disease can be corrected by gene editing protocols in which endogenous cellular DNA repair mechanisms make use of an exogenous donor template to repair a double-stranded break (DSB) created by the Cas9 endonuclease at or near to the site of the mutation. ssODNs are attractive templates for HDR because they can readily be synthesized with a specified sequence and length, and avoid production costs and toxicity inherent in viral DNA templates^1^. Although oligonucleotide synthesis is a mature technology, the quality of the synthesis product is routinely assessed only by liquid chromatography, gel electrophoresis, and mass spectrometry, with sequence fidelity not directly confirmed. The sequence fidelity of chemically synthesized oligonucleotides has been characterized in a few previous studies. Filges et al. showed that deletions predominate over substitutions, with error profiles that vary by manufacturer, purity grade, batch, and sequence context^2^. Further studies demonstrated that substitution errors, particularly G-to-A transitions, are strongly influenced by the capping step of phosphoramidite synthesis, and proposed modifications to synthesis chemistry that reduce their frequency^3,4^. Collectively, these studies establish that synthesis errors in ssODNs are reproducible and sequence-context-dependent but share the critical limitation that their sequencing approaches required oligonucleotides specifically designed to enable library construction. These methods are therefore not applicable to the characterization of ssODNs whose sequences are fixed by therapeutic design, as is the case in clinical gene editing protocols. Furthermore, none of these studies addressed whether synthesis errors are propagated into the genome when an ssODN is used as a donor template for HDR, leaving the consequences of synthesis errors for therapeutic applications uncharacterized. A sensitive and broadly applicable means of assessing sequence fidelity is needed.

We developed a gene editing protocol to correct the mutation in the human β-globin gene (*HBB*) that causes Sickle Cell Disease (SCD)^5,6^. Cas9 is directed by a single guide RNA (sgRNA) to introduce a double-stranded break in *HBB* 18 base pairs (bp) from the SCD mutation in autologous hematopoietic stem and progenitor cells (HSPCs). The 168 bp ssODN has a sequence complementary to the region of *HBB* encompassing the SCD mutation, restoring an adenine base to correct the SCD mutation when used as a donor template for HDR. Clinical application of this protocol, and of any protocol aiming to correct a mutation using an ssODN, requires a tractable means to validate the identity and sequence of the ssODN and to understand what, if any, sequence changes might accompany correction of the mutation. We have addressed this need by creating deep sequencing libraries from oligonucleotides and corroborating apparent sequence errors by showing that they are propagated into the genome by HDR.

We used three different vendors to manufacture the ssODN. While the purity of an ssODN was determined by the manufacturer using Ion Pairing HPLC and SDS-PAGE, and its identity was evaluated by liquid chromatography/electrospray ionization mass spectrometry, neither method confirms the specific nucleotide sequence of an ssODN or the absence of synthesis errors. To establish a method to quantify ssODN sequence fidelity and determine the best practices, we performed very deep sequencing of the ssODNs. We prepared sequencing libraries (**Figure 1**) using two different library preparation kits that are optimized for ligating adapters to single-stranded DNA: the SRSLY PicoPlus kit from Claret Bioscience which uses unique molecular identifiers (UMIs) to combat PCR bias, and the xGen kit from IDT which does not use UMIs. Each ssODN was sequenced with a minimum of three million 150bp paired-end reads. Following alignment to our reference ssODN sequence and read deduplication using the UMIs (for the SRSLY kit), we achieved an average minimum depth of 150,000 aligned reads (**Supplementary Figure 1**).

**Figure 1.**
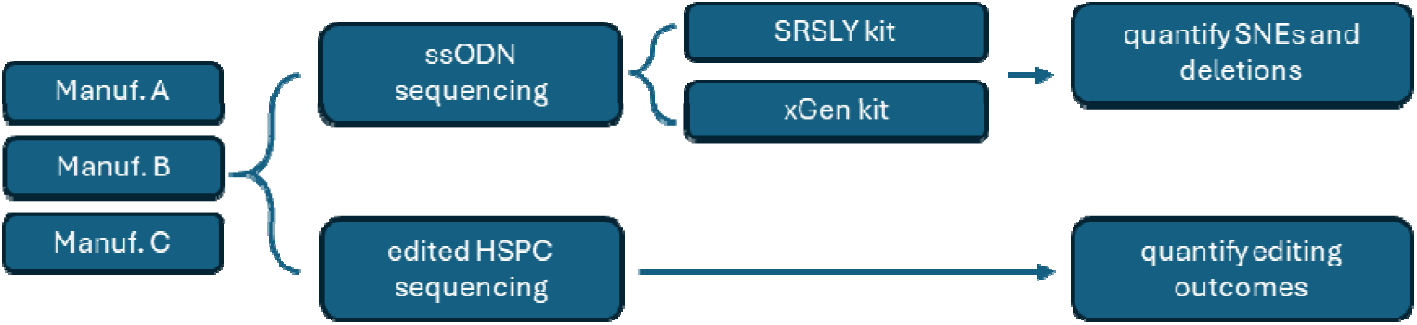
Schematic of sequencing strategy for three ssODNs, and of the SCD locus following editing of HSPC with each of the ssODNs. SNEs and deletions in the ssODNs were compared to editing outcomes.

Analysis of the data aligned to the ssODN reference revealed single nucleotide errors (SNEs) and short deletions at a much higher frequency than would be expected due to sequencing errors or PCR bias alone. Notably, the frequency of both types of error varied dramatically by manufacturer, with the SSODN from manufacturer A typically having over two-fold more errors than the ssODNs from manufacturers B and C. SNEs were found to occur more frequently at specific positions within the ssODN sequence across all three vendor-sourced ssODNs (**Figure 2A, Supplementary Figure 2A**), while small deletions (typically 1-2bp in length) were more uniformly distributed across the sequence (**Figure 2B, Supplementary Figure 2B**). The rate of errors at individual positions was as high as 9% for SNEs and 3% for deletions, with the highest abundance seen in the ssODN from manufacturer A for SNEs and manufacturer C for deletions. The SNEs do not involve specific nucleotide conversion: for example, a site where a C nucleotide was specified might be modified primarily to a T, but also have changes to G or A.

**Figure 2.**
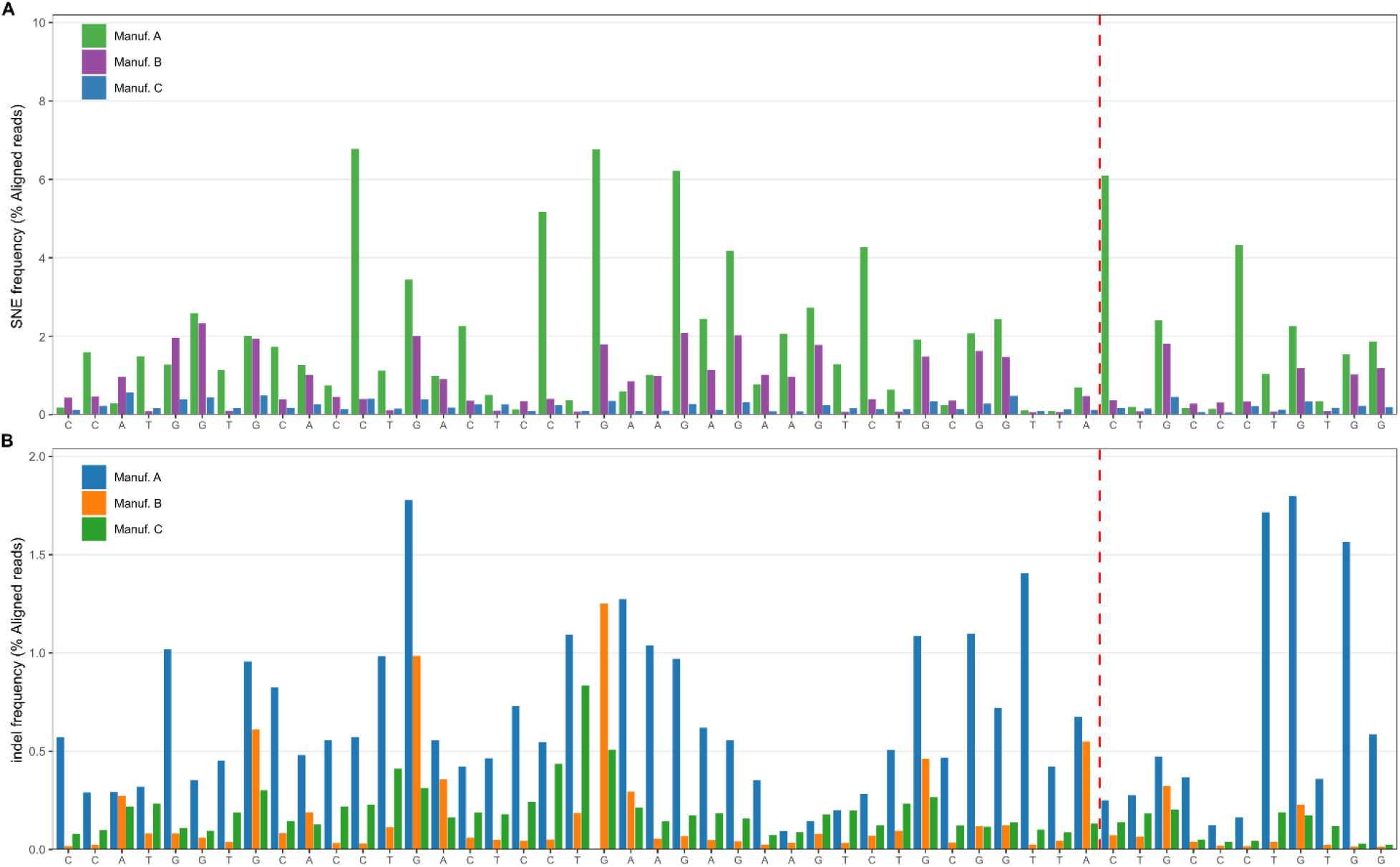
SNE (**A**) and indel (**B**) frequency in the three ssODNs, determined by direct sequencing. The x-axis shows a 50-bp window containing the Cas9 cleavage site specified by the guide RNA (vertical dashed red line) is shown, the PAM motif, and the SCD mutation; similar SNE levels were observed outside this region (see Supplementary Figure 2). The y-axis shows SNE frequency calculated as the percentage of aligned reads that contains an error.

To investigate whether specific nucleotide identities or local sequence contexts are associated with elevated error rates, we applied a binomial enrichment test to identify mononucleotides and dinucleotides over-represented among the top 20% highest-error positions in the ssODNs from each manufacturer. Relative to the background base composition of the ssODN, G was significantly over-represented at high-error positions in all manufacturers (**Figure 3A**, p_adj_ < 0.05, fold over-representation between 2-3 depending on manufacturer); C-containing positions were also over-represented in Manufacturer A, although to a lower degree. To determine if the base preceding the error position during synthesis predicted the occurrence of an error, we examined the frequency of dinucleotides formed by error positions and the base immediately 3’ to them. We found that G-containing error positions tended to be preceded by A or T (**Figure 3B**, p_adj_ < 0.05, fold over-representation between 3-4 depending on manufacturer); in addition, C was also found to precede a G in Manufacturers B and C, and CT dinucleotides were more than 4-fold over-represented in Manufacturer A. These specificities of synthesis error sequence contexts are broadly consistent with previous studies^2–4^, and indicate that error-prone positions share identifiable sequence features, and that the relationship between sequence context and error rate differs among manufacturers, potentially reflecting differences in synthesis chemistry or post-synthesis processing.

**Figure 3.**
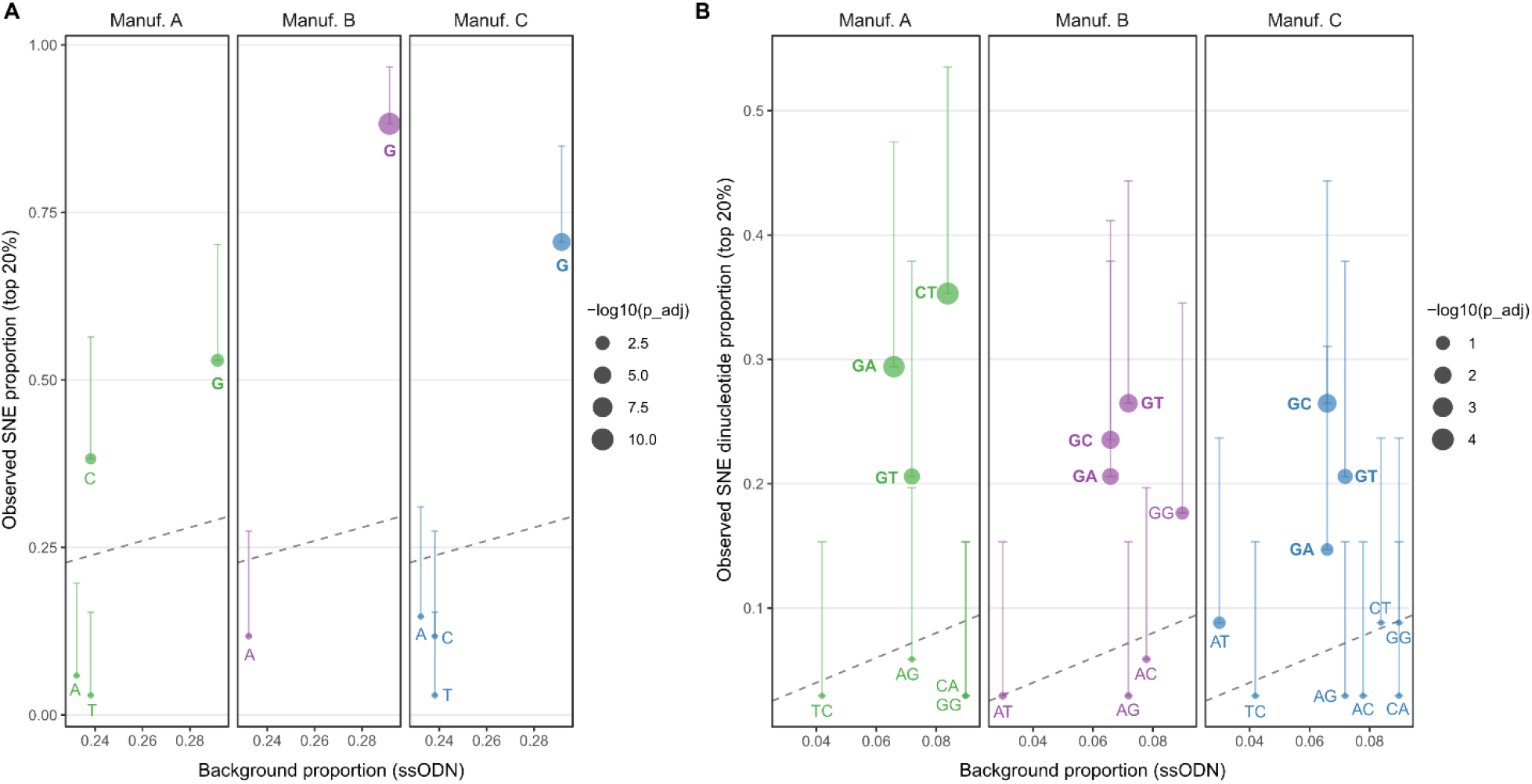
Nucleotide context enrichment at high-error positions in the ssODN. (**A**) Mononucleotide fold enrichment: individual bases significantly over-represented (BH-adjusted binomial test, p_adj_ < 0.05) among the top 20% highest-error positions relative to background base composition of the ssODN. (**B**) Dinucleotide fold enrichment: dinucleotide contexts significantly over-represented at high-error positions. Points show fold enrichment (observed/background proportion); horizontal bars extend to the lower 95% confidence bound. Manufacturers are distinguished by color. The dashed diagonal line marks significance at p_adj_=0.05

To directly test whether ssODN synthesis errors are propagated into the genome by HDR and exclude that the observed SNEs and deletions in the ssODNs were the result of PCR or sequencing errors, we compared SNE profiles from direct ssODN sequencing with those from amplicon sequencing of edited HSPCs. We reasoned that when an ssODN is used in a gene editing protocol, errors in its sequence will be incorporated into the genome by HDR. However, while SNEs are present throughout the 168bp ssODN, only those present in the region where the ssODN is used as a template for HDR (HDR region) will be incorporated into the genome. Thus, true ssODN synthesis errors will produce an excess of sequence variants in edited genomic DNA only in the vicinity of the Cas9 cleavage site, where HDR occurs. In contrast, errors that result from the ssODN sequencing process will not be present in the genomic DNA following editing. When ssODNs with high rates of errors are used as a template for HDR, we observe that the errors are incorporated into the genomic DNA of edited cells. We analyzed targeted amplicon sequencing data generated from the region surrounding the SCD mutation in both SCD and healthy donor HSPCs that had been edited with our protocol using ssODNs from the three vendors (**Figure 1**). Amplicons derived from HBB alleles that were subjected to HDR can be identified by the presence of sequence changes programmed into the ssODN: an alteration in the PAM motif, reversion of the sickle mutation to wild type, and a neutral substitution in the base preceding the sickle mutation that allows identification of HDR in wild type cells ^5,6^. The sequenced amplicon encompasses a 350bp window surrounding the Cas9 cut site and the entire sequence of the ssODN. **Figure 4A** overlays SNE frequency at each position in the ssODN with SNE frequency in HDR-derived genomic reads from multiple independently edited HSPC samples. Strikingly, high-frequency SNE positions in the ssODN (colored red, indicating the top quintile error rate) correspond to positions with elevated SNE frequency in the genomic amplicons, while positions with low ssODN error rates (green, bottom three quintiles) show correspondingly low SNE frequencies in the genomic DNA. This positional concordance is consistent with direct templating of genomic sequence by the error-containing ssODN during HDR. Furthermore, SNEs are incorporated into the HSPCs at rates that recapitulate the variation by manufacturer seen in the direct ssODN sequencing (**Supplementary Figure 3**).

**Figure 4.**
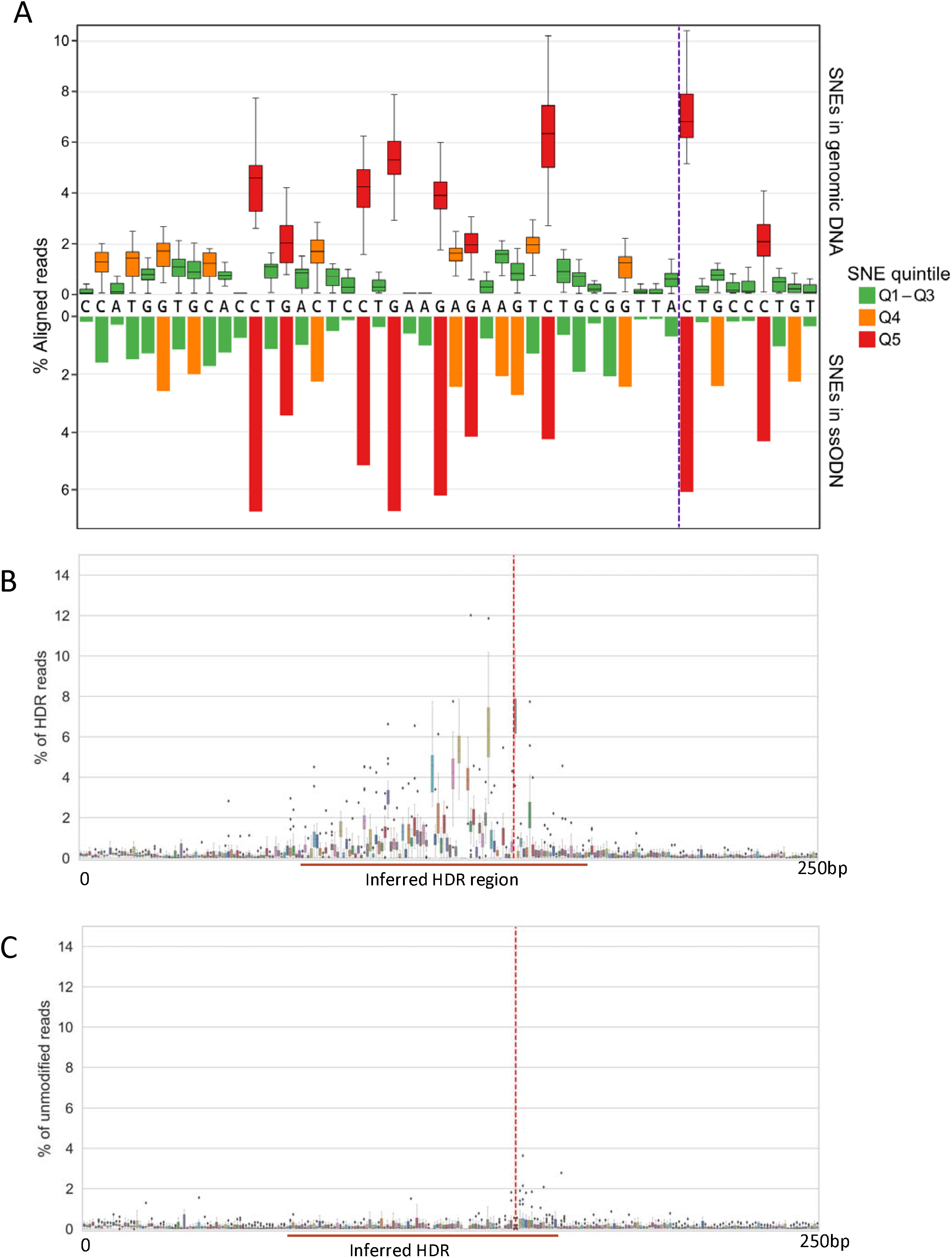
SNE frequency in genomic DNA from HSPCs edited with ssODN from Manufacturer A. (**A**) Comparison of SNEs in ssODN and edited genomic DNA. The x-axis shows the region containing the Cas9 cleavage site (marked by the vertical purple line), the PAM motif, and the SCD mutation. The y-axis shows SNE frequency calculated as the percentage of aligned reads that contains an error. Error frequencies are colored by quintile: green indicates positions in the bottom three quintiles (low frequency), orange the fourth quintile, and red the fifth quintile (highest frequency errors). *Upper panel*: SNEs in genomic DNA isolated from multiple edited HSPC samples using ssODN from manufacturer A were quantified and displayed as boxplot. *Lower panel*: SNEs detected by direct sequencing of ssODN from manufacturer A, shown as a reference for template-associated errors. (**B**) SNE frequency in HDR-derived reads from genomic DNA (HDR-derived reads are defined by editing of the PAM site). A 250-base window containing the cleavage site is shown, with the inferred HDR template region labeled. The Cas9 cleavage site is indicated by the vertical dashed red line. The HDR region spans approximately 50-75bp to the left of the cleavage site to 10bp to its right. Although the plot encompasses the full 168-bp ssODN, errors are incorporated into a smaller region of ~60–85 bp centered on the cleavage site, consistent with the boundaries of the HDR template. The absence of SNEs outside this region strongly suggests that observed errors derive from the ssODN template. (**C**) In amplicons derived from unedited reads, SNEs were rarely observed within or outside the HDR region. A small excess of SNEs proximal to the cleavage site likely reflects HDR events that initiated at the cleavage site but did not extend to the PAM site, precluding definitive classification as HDR-derived reads.

SNEs in the three ssODN were consistently observed at the same positions in the amplicon sequencing data at a frequency roughly correlated with SNE frequency in the ssODNs. Outside of the region where the ssODN was used as a template for HDR, we see errors at a rate that would be expected from PCR bias or sequencing errors, and comparable to the rate observed in reads from unedited samples (**Figure 4B**). Deletions, which occurred more uniformly across the length of the ssODN, were evenly distributed only in the HDR region, and not in the surrounding sequence (**Supplementary Figure 4**). Furthermore, the ssODN SNEs were not observed in unmodified reads (alleles that lacked base changes at the PAM and the site of the sickle mutation) (**Figure 4C**).

The presence of SNEs in the ssODN also provides valuable information on the extent and polarity of HDR in our protocol. We noted that the frequency of SNE propagation by HDR (determined as the frequency of SNEs in the genome amplicon sequence) was consistent with HDR initiating at the Cas9 cleavage site, proceeding largely asymmetrically (to the left of the cleavage site in **Figure 4A**), and attenuating with distance from the cleavage site. Thus, ssODN SNEs can be used as markers of the HDR conversion tract to define the genomic region where the ssODN was used as a template for HDR at highest frequency: in our system, this was the genomic interval roughly 50-75bp upstream and 10 bp downstream of the Cas9 cleavage site.

In sequenced alleles that had undergone HDR, rates of incorporation of ssODN synthesis errors into genomic DNA occurred at a frequency as high as 12% for SNEs and up to 3% for deletions at individual positions. Incorporation of SNEs was observed with ssODNs from all three manufacturers at a rate proportional to the SNE frequency in the respective ssODNs (**Figure 4A, Supplementary Figure 3**). Deletions in the ssODNs were also incorporated into the genomic DNA by HDR, but at a lower frequency than the SNEs (**Supplementary Figure 4**). Significantly, 90% of the deletions seen were 1 or 2bp frame-shift deletions: although they occur at a lower frequency than SNEs, they would have a more consequential effect on β-globin production.

This investigation of edited cell products thus corroborates the direct sequencing of ssODNs and highlights the variation in synthesis errors among manufacturers (**Figure 2, Supplementary Figure 2**). The basis of the ssODN synthesis errors was not apparent to any of the manufacturers, all of whom used the standard chemical synthesis by coupling of phosphoramidite analogues and were unaware of the presence of sequence errors in the synthesis products.

Synthesis errors in ssODN templates for HDR-mediated gene editing have implications for both pre-clinical and clinical development. The method we have used improves on previous efforts^2–4^ by allowing investigators to establish the extent of sequence errors in any ssODN using standard reagents. The enrichment of errors at C and G positions is consistent with known challenges in phosphoramidite-based synthesis at consecutive or isolated G and C residues and may help guide future improvements to synthesis protocols. None of the prior studies determined whether synthesis errors are faithfully propagated into the genome by HDR; here, we demonstrate that SNEs and deletions present in the donor template are incorporated into edited alleles. The quantitative relationship between ssODN SNE frequency and genomic SNE frequency implies that improvements in ssODN synthesis quality will translate directly into reduced error burden in edited alleles. Critically, these errors are invisible to standard QC methods (HPLC, mass spectrometry), which assess overall purity and molecular weight but cannot resolve single-nucleotide compositional errors at specific positions. The data therefore argue that the sequencing approach described here should be a prerequisite for regulatory submissions for clinical gene editing programs, both to characterize the error profile of any given batch and to enable informed risk assessment of the amino acid consequences of the most prevalent synthesis errors.

The effects of SNEs and deletions on editing outcomes will vary depending on the specific gene and the type of sequence modified: errors may have more impact in coding regions, but the effects of specific errors in transcriptional control elements are less predictable. In the example of our protocol, available clinical data predict that amino acid changes caused by SNEs in the edited region of *HBB* are likely to create benign ß-globin variants, while frameshift deletions will create potentially deleterious ß-thalassemia-like alleles^7–11^.

## AUTHORS CONTRIBUTIONS

SW, DIKM, and DB conceived the study; ZR, SJH, MN, and NK performed the experimental work; SW and DB analyzed the data; SW prepared all the figures; SW, DBK, MCW, DIKM, and DB interpreted the results and wrote the manuscript. All authors reviewed the manuscript.

## AKNOWLEDGMENTS

Support to this study was provided to Mark Walters by CIRM (TRAN1-09292, CLIN1-11497, and CLIN2-11722) and NIH Cure Sickle Cell/NHLBI OT2 HL151319 and OT3HL147741.

## DATA AVAILABILITY

The sequence data used in this study are available in the NCBI SRA archive as BioProject PRJNA1281709.

## METHODS

### ssODN library preparation and sequencing

ssODN sequencing libraries were constructed using the Claret Bioscience SRSLY ssODN library preparation kit and IDT’s xGen RNA Library Preparation Kit following the manufacturer’s instructions. Libraries were sequenced using Illumina 150bp paired end reads to a minimum depth of 3M reads.

Sequenced reads were trimmed and merged to single reads using bbmerge, then aligned to the reference ssODN using bowtie2. Where UMI’s were used, reads were deduplicated using UMI-tools. SNEs, insertions and deletions were then quantified using samtools and custom Python scripts (see flow chart, Supp Fig X).

### Editing of HSPCs

Editing of HSPCs with the three ssODNs was carried out as described in Dewitt et al. 2016 and Magis et al. 2022

### Amplicon Sequencing

HBB amplicon generation and sequencing were carried out as described in Dewitt et al. 2016 and Magis et al. 2022

### Data Analysis

Analysis of sequencing data for direct ssODN sequencing was performed as described in Supplementary Figure 1. Rates of incorporation into the genomic DNA were calculated using the amplicon sequencing data. The mismatches and deletions from the read alignment were tallied for all reads to get the rate of SNEs for an individual sample. SNE frequency across samples was then compiled together for each ssODN and plotted as boxplots. Each box represents and individual position around the cut site in the sequenced amplicon. Deletions were counted on the first (left hand) nucleotide position of the deletion so that they were counted just once per deletion (versus in each position that the deletion spans).

### Nucleotide context enrichment analysis

For each manufacturer, positions were ranked by SNE error rate and the top 20% were designated high-error positions. Mononucleotide and dinucleotide frequencies among high-error positions were compared to their background frequencies in the full ssODN sequence using one-sided binomial tests (alternative = “greater”). P-values were adjusted for multiple testing using the Benjamini-Hochberg procedure. Fold enrichment was calculated as the ratio of observed to background proportion; confidence intervals are derived from the lower bound of the exact binomial confidence interval for the observed proportion, scaled by the background proportion. All analyses were performed in R using base stats and the tidyverse suite.

---

**Supplementary Figure 1:**
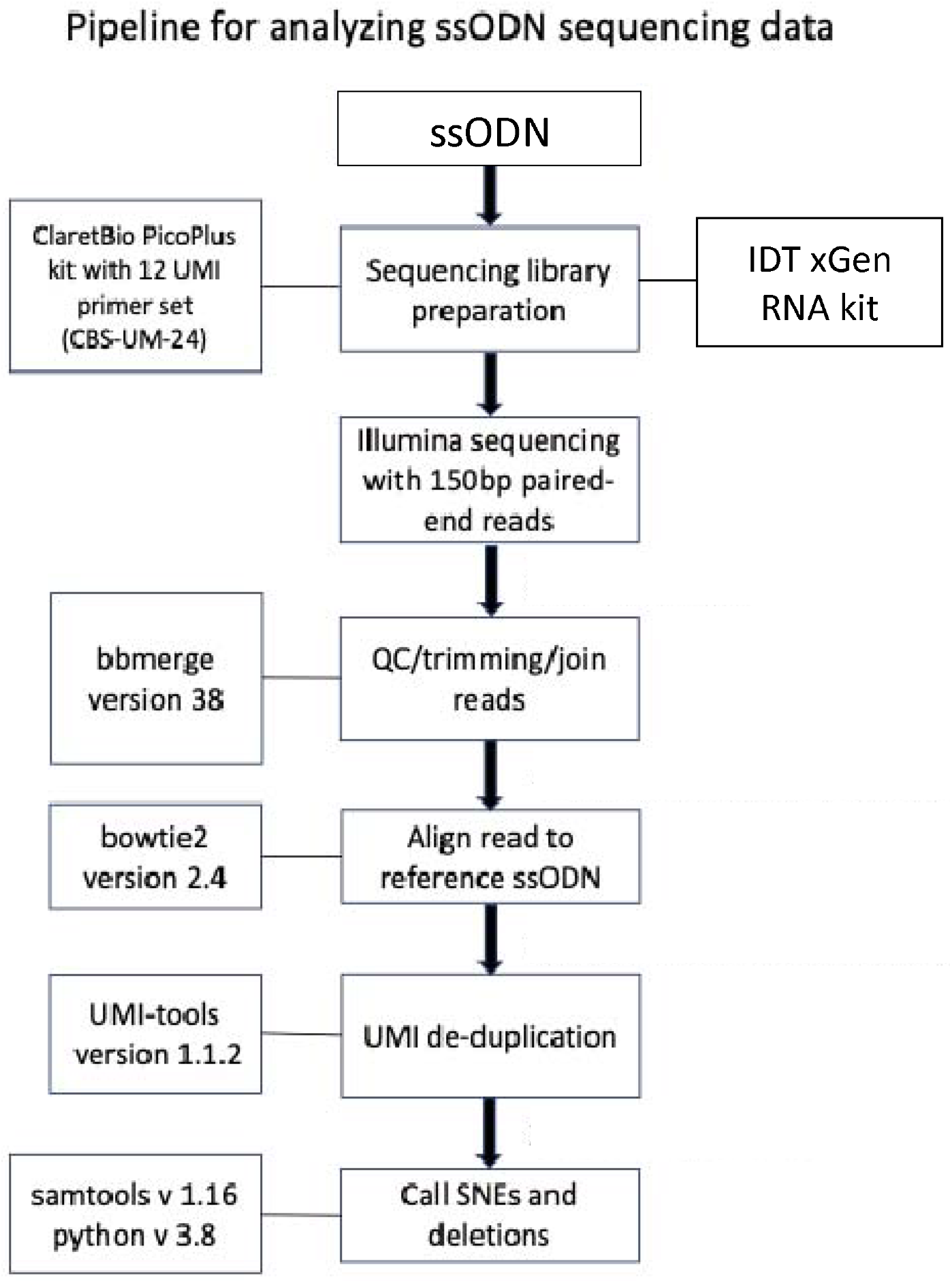
Schematic of analysis pipeline.

**Supplementary Figure 2:**
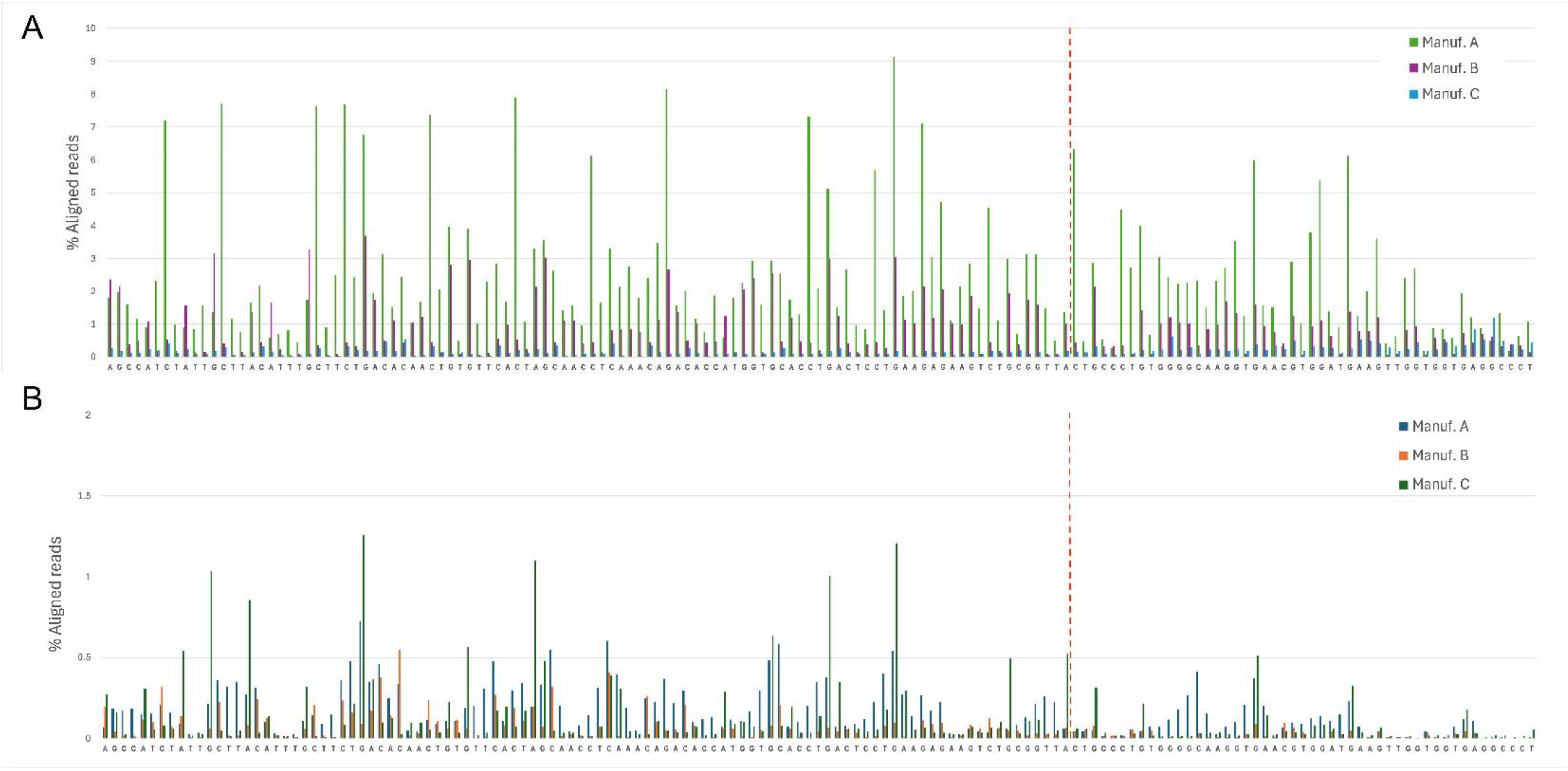
SNE (**A**) and indel (**B**) frequency in the three ssODNs, determined by direct sequencing. The full length ssODN is shown, with 8 bases pruned from each end to eliminate library preparation artifacts that appear at the ends. Cas9 cleavage location is marked by a vertical dashed red line (see Figure 2 for a detailed view of the 50-base region surrounding the Cas 9 cut site). Error frequency (SNE or indel) was calculated as the fraction of aligned reads with an error at a given position.

**Supplementary Figure 3:**
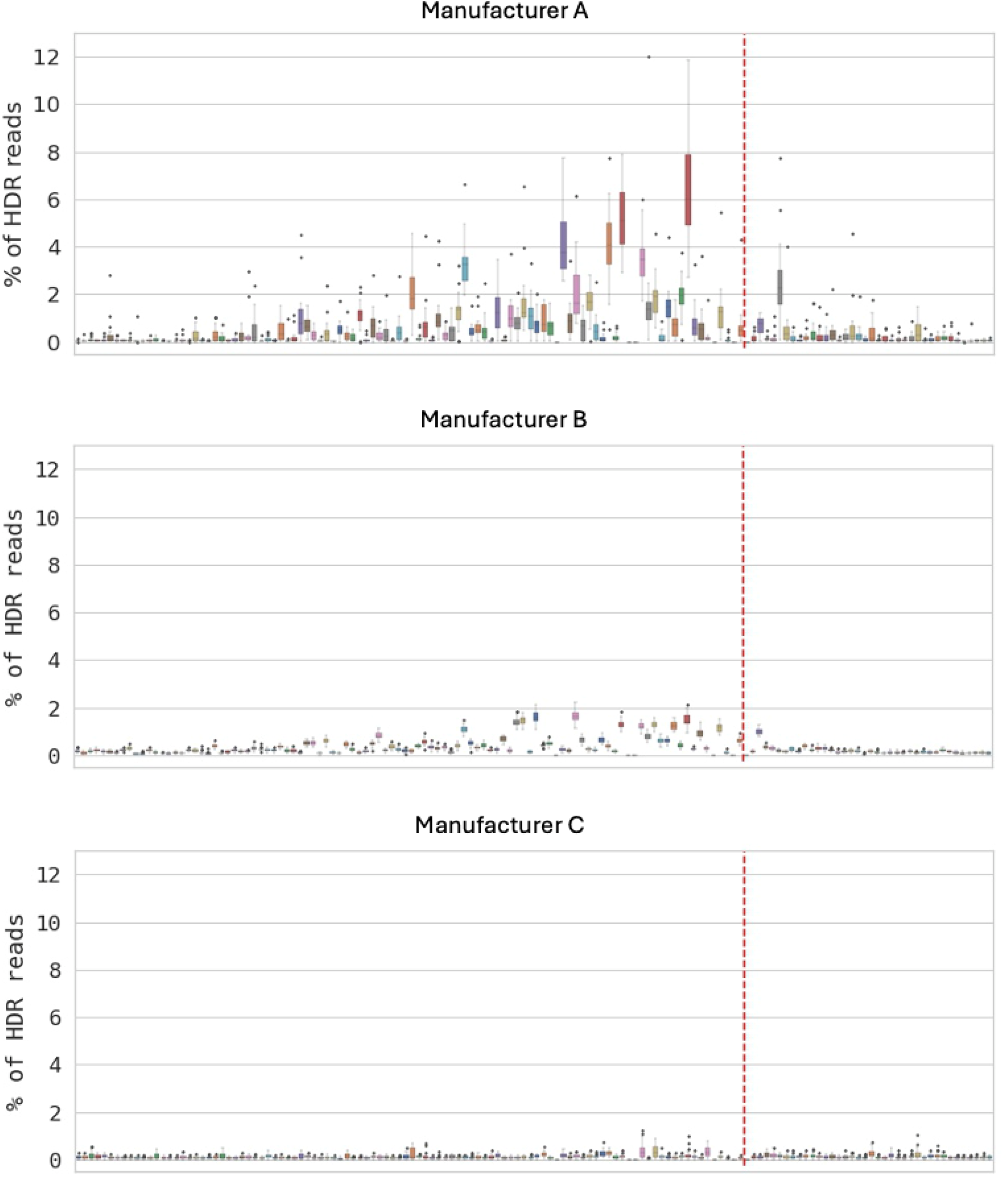
SNEs in genomic DNA from multiple edited HSPC samples using ssODNs from three different manufacturers were quantified as a percent of HDR reads and plotted. The dashed red line represents the Cas9 cleavage site.

**Supplementary Figure 4:**
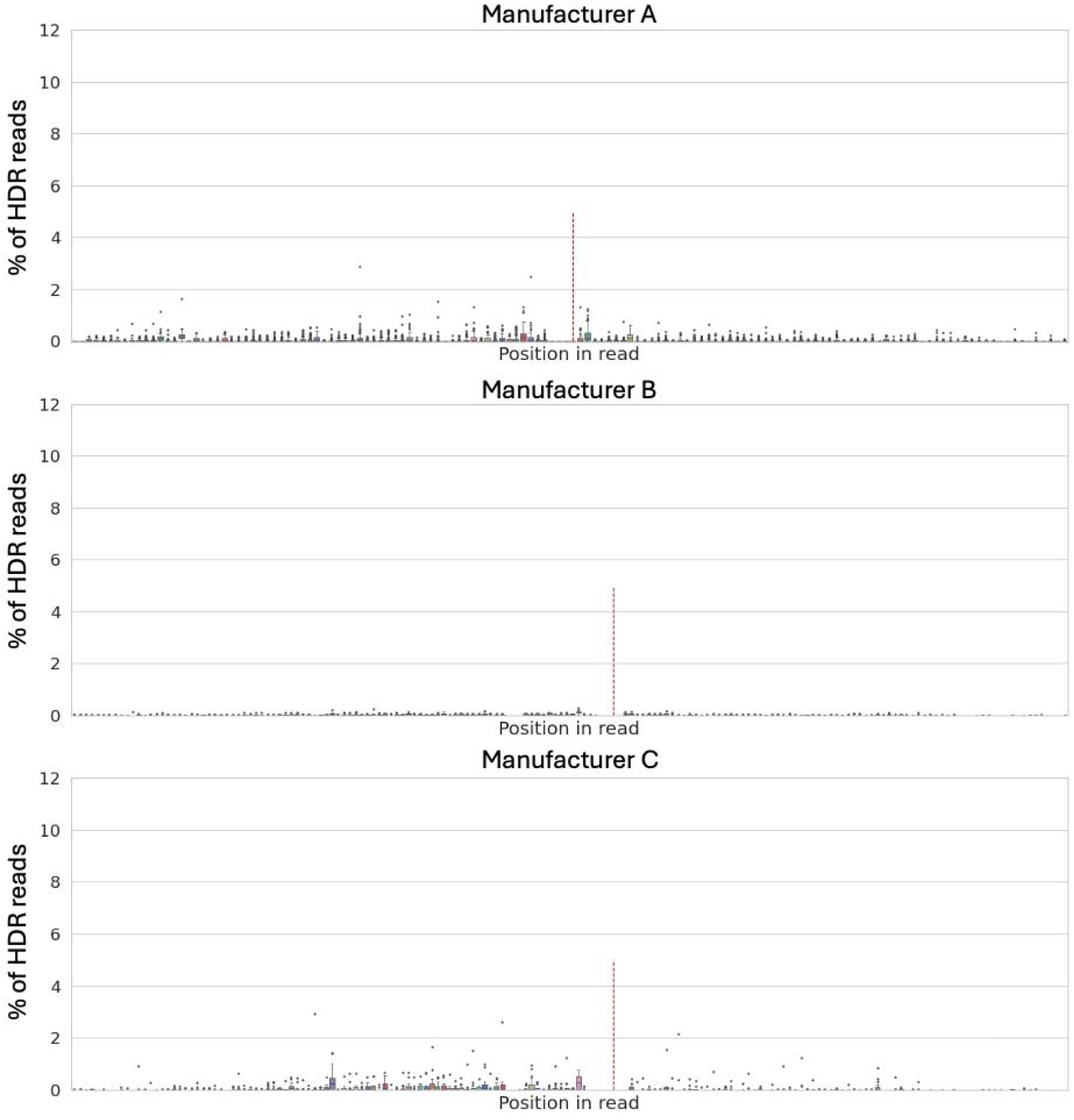
Deletions in genomic DNA from multiple HSPC samples edited using ssODNs from the three different manufacturers, quantified as a percent of HDR reads and plotted. Dashed red line represents cleavage site. Deletions are not seen directly adjacent to the cleavage site because reads with deletions at the cleavage site were counted as NHEJ reads.

